# Using predictive machine learning models for drug response simulation by calibrating patient-specific pathway signatures

**DOI:** 10.1101/2020.12.06.413435

**Authors:** Sepehr Golriz Khatami, Sarah Mubeen, Vinay Srinivas Bharadhwaj, Alpha Tom Kodamullil, Martin Hofmann-Apitius, Daniel Domingo-Fernández

## Abstract

The utility of pathway signatures lies in their capability to determine whether a specific pathway or biological process is dysregulated in a given patient. These signatures have been widely used in machine learning (ML) methods for a variety of applications including precision medicine, drug repurposing, and drug discovery. In this work, we leverage highly predictive ML models for drug response simulation in individual patients by calibrating the pathway activity scores of disease samples. Using these ML models and a novel scoring algorithm to modify the signatures of patients, we evaluate whether a given sample that was formerly classified as diseased, could be predicted as normal following drug treatment simulation. We then use this technique as a proxy for the identification of potential drug candidates. Furthermore, we demonstrate the ability of our methodology to successfully identify approved and clinically investigated drugs for three different cancers. We also show how this approach can deconvolute a drugs’ mechanism of action and propose combination therapies. Taken together, our methodology could be promising to support clinical decision-making in personalized medicine by simulating a drugs’ effect on a given patient.

## 1. Introduction

Within the biomedical field, the application of numerous technologies and machine learning (ML) and artificial intelligence (AI) methods have enormous potential for the development of personalized therapies (Pai *et al*., 2019), drug repurposing (Zhao and So, 2019), and drug discovery (Réda *et al.,* 2020). The data exploited by these methods can comprise multiple modalities including imaging data (Liu *et al.*, 2014), chemical structure information (Hirohara *et al*., 2018), and natural language data (Castro *et al*., 2017). However, the widespread availability of transcriptomics data (e.g., RNA-Sequencing (RNA-Seq), microarrays, etc.) along with its capacity to provide a comprehensive overview of biological systems have made this particular modality a popular choice for various computational methods. Although this modality can reveal both molecular signatures as well as phenotypic changes that occur in altered states, pathway analyses are often performed to map measured transcripts to the pathway level due to high dimensionality and correlations present in transcriptomics datasets (Su *et al*., 2009; Lim *et al.*, 2020). This transformation facilitates the training of ML/AI models by reducing dimensional complexity whilst enhancing interpretive power (Reimand *et al*., 2019). However, such a transformation implicates the use of prior pathway knowledge (Perscheid, 2020) from databases such as KEGG (Kanehisa *et al.*, 2017) and Reactome (Jassal *et al.*, 2020) (Nguyen *et al.,* 2019).

The transformation of data from the transcriptomics to the pathway level can be used to generate pathway features (i.e., sets of genes involved in a given pathway that are coordinately up or down-regulated), the latter of which have broad applications in drug discovery and drug response prediction (Adam *et al*., 2020). For instance, Peyvandipour *et al*. (2018), Saberian *et al*. (2019), and Emon *et al*. (2020) exploited the concept of anti-similarity between drugs and disease-specific pathway signatures to identify therapeutic candidate drugs that can potentially revert disease pathophysiology. Furthermore, Ammad-ud-din *et al.* (2016) show how pathway signatures derived from cell lines using kernelized Bayesian matrix factorization can be used for drug response prediction.

Alternatively, other methods can generate individualized pathway features from a population of patients or cell lines (Amadoz *et al*. 2019). These features, or pathway activity scores, can subsequently be used for several downstream ML applications including classification tasks and survival prediction (Lim *et al*., 2020; Mubeen *et al.*, 2019). Additionally, Wang *et al*. (2019) showed how ML models can be used to predict drug response using pathway activity scores derived from cell lines. Furthermore, another example from Esteban-Medina *et al*. (2019) demonstrated how modelling individualized pathway activity scores from Falconi anemia patients can reveal potential targets for therapeutic intervention.

While these methods have shown how pathway signatures can be used for drug discovery and drug response prediction, existing methods thus far fail to account for two important factors. Firstly, as the response triggered by a drug in a given patient may differ if administered in another, these methods should account for patient heterogeneity which is crucial in designing individualized therapies. Secondly, specific indications may be improved or corrected by a drug combination approach or through the administration of multi-target drugs.

In this work, we present a novel methodology that exploits the predictive power of ML models to simulate drug response by calibrating pathway signatures of patients. We first trained an ML model (i.e., elastic net penalized logistic regression model) to discriminate between disease samples and controls based on sample-specific pathway activity scores. Next, we simulate drug responses in patients using a scoring algorithm that modifies a patient’s pathway signatures using existing knowledge on drug-target interactions. We hypothesize that promising drug candidates for a given condition would modify pathway activity scores of patients in such a way that they closely resemble scores of controls. Thus, using the previously trained ML model, we then evaluate whether patients with modified pathway scores are now classified as normal as a proxy for promising drug candidates. We demonstrate the scalability and generalizability of our methodology by simulating over one thousand drugs on three cancer datasets. Furthermore, we show how our methodology is able to recover a large proportion of clinically investigated drugs on these three indications. Finally, we show how the most relevant pathways identified by our methodology can be used to better understand the biology pertaining to a given condition and propose new targets.

## 2. Methodology

The initial step of our methodology consists of generating patient-specific features that can be used for model training. Although in this work, we employed pathway activity scores **(Subsection 2.2)**, other features could also be used for the same purpose. Using these scores, we trained an ML model **(Subsection 2.3; Figure 1a)**that can accurately discriminate between sample classes (e.g., disease vs normal). Next, we developed a scoring algorithm aimed to simulate the effect of a drug intervention at the pathway-level by modifying the pathway activity scores of disease samples **(Subsection 2.4; Figure 1b)**. The final step of our methodology uses the modified pathway activity scores as an input in the trained model to assess whether samples that were previously classified as “diseased” could now be classified as “normal” as a proxy for drug candidates **(Subsection 2.5; Figure 1c)**. Finally, in the last three sections, we present the validation datasets used for our case scenario **(Subsection 2.6; Figure 1d-f),**compare our methodology against six equivalent approaches **(Subsection 2.7)**, and provide details on implementation and reproducibility **(Subsection 2.8)**.

**Figure 1.**
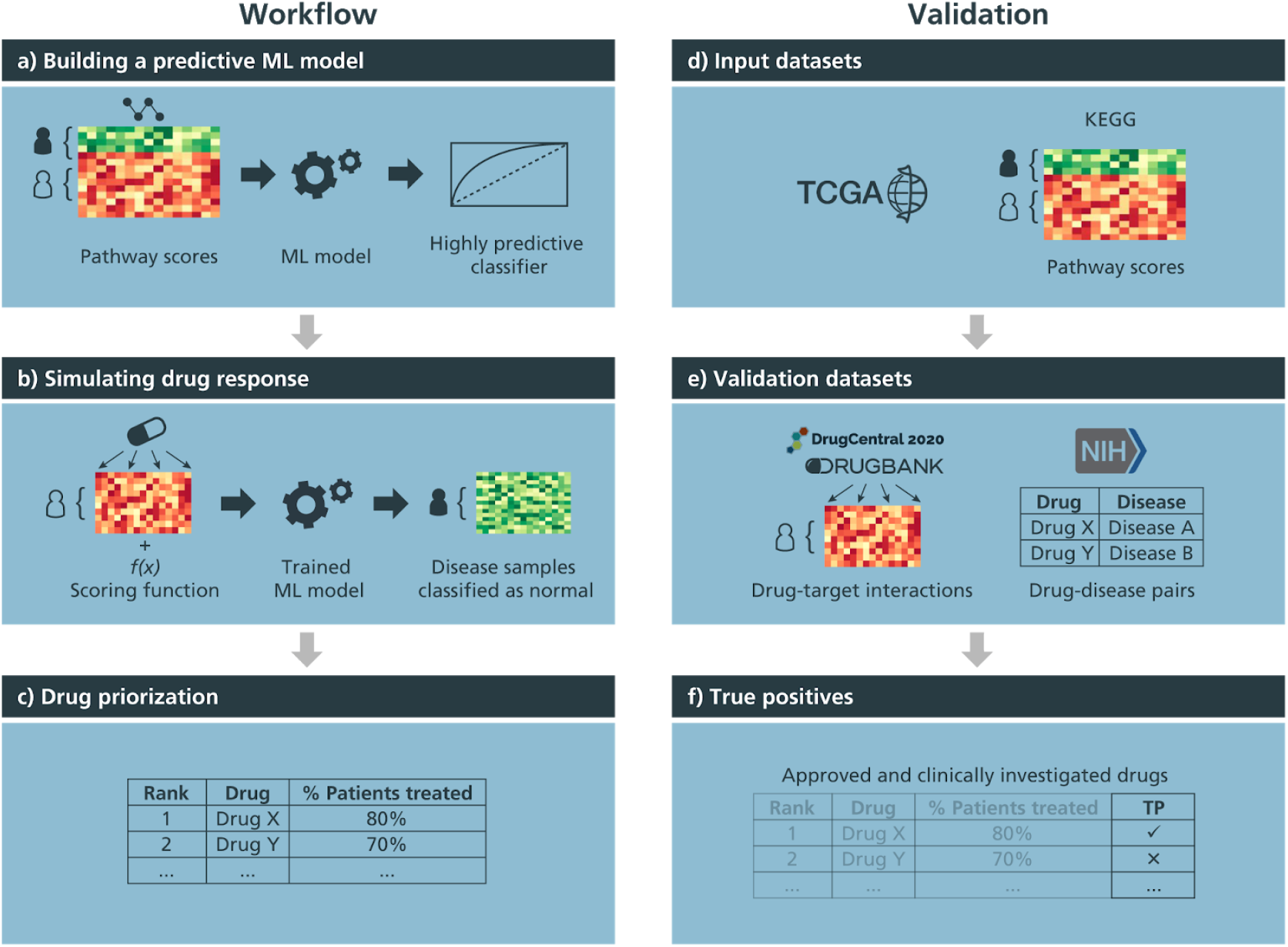
Conceptual overview of the methodology and method validation. **a)** Pathway activity scores are used to train a highly predictive ML model that differentiates between normal and disease samples, labelled green and red on the heatmap, respectively. **b)** Next, pathway scores of disease samples are modified by using drug-target information and applying a scoring algorithm that simulates the effect of a given drug at the pathway-level. The trained ML classifier was then used to evaluate whether the modified disease samples that were previously classified as “diseased” could now be classified as “normal”. **c)** Finally, we used the proportion of disease samples now classified as normal as a proxy to identify candidate drugs, propose combination therapies, and pinpoint new targets. **d)** To validate the methodology, we first performed ssGSEA using pathways from KEGG and the BRCA-, LIHC- and PRAD-TCGA datasets to acquire sample-wise pathway activity scores. **e)** Next, we obtained known drug-target interactions from DrugBank and DrugCentral and drug-disease pairs (i.e., FDA-approved drugs and drugs under clinical trials for a given condition) from Clinicaltrials.gov and FDA-approved drugs, of which, the latter two were used as a ground-truth list of true positives (TP). **f)** To simulate drug treatments of patients from the aforementioned TCGA datasets using their pathway activity scores (i.e., Figure 1d), we applied the methodology described in Fig. 1a-c to acquire a ranking of drugs based on the proportion of disease samples that were treated. Finally, we identified the proportion of drugs ranked by our methodology that were true positives for the three TCGA datasets and compared this proportion to random chance.

### 2.1. Datasets

Datasets from The Cancer Genome Atlas (TCGA) (Weinstein et *al.*, 2013) were retrieved from the Genomic Data Commons (GDC; https://gdc.cancer.gov) portal through the R/Bioconductor package, TCGAbiolinks (version 2.16.3; Colaprico *et al.*, 2015) on 04-08-2020 **(Figure 1d)**. Gene expression data from RNA-Seq was quantified using the HTSeq and raw read counts were normalized using Fragments Per Kilobase of transcript per Million mapped reads upper quartile (FPKM-UQ). Gene identifiers were mapped to HUGO Gene Nomenclature Committee (HGNC) symbols where possible. The datasets downloaded include The Cancer Genome Atlas Breast Invasive Carcinoma (TCGA-BRCA), The Cancer Genome Atlas, Prostate Adenocarcinoma (TCGA-PRAD) and The Cancer Genome Atlas Liver Hepatocellular Carcinoma (TCGA-LIHC) **(Supplementary Table 1)**.

### 2.2. Calculating individualized pathway activity scores

We used single-sample GSEA (ssGSEA) (Barbie *et al.*, 2009), a commonly used tool to generate patient-specific pathway activity scores. Normalized gene expression (FPKM-UQ) and pathway definitions (i.e., gene sets) were provided as input and were converted to scores through ssGSEA. As a reference database, we used 337 pathways from KEGG (downloaded on 01-04-2020) as it is the most widely used pathway database and a standard for the most commonly used pathway activity scoring methods (Amadoz *et al.*, 2019) **(Figure 1d)**.

### 2.3. Building a predictive classifier

Patient-specific pathway activity scores generated by ssGSEA were used to generate a ML classifier to distinguish between normal and tumor sample labels for each of the three datasets. The classification was conducted using an elastic net penalized logistic regression model (Zou and Trevor, 2005) as regularized models have been shown to achieve superior predictions and better generalize to unseen data (Wang *et al*., 2019). Furthermore, we previously used this ML model on the same TCGA datasets (Mubeen *et al*., 2019), yielding AUC-ROC and AUC-PR values close to 1 **(Supplementary Figure 1)**. Prediction performance was evaluated via 10 times repeated 10-fold stratified cross-validation and tuning of elastic net hyper-parameters (i.e., *l*_1_, *l*_2_ regularization parameters) *via* grid search was performed within the cross-validation loop to avoid over-optimism (Molinaro *et al*., 2005).

### 2.4. Scoring algorithm

To modify the pathway activity scores for disease samples, we developed a scoring algorithm to replicate the effect of a drug at the pathway-level. The scoring algorithm exploits interactions from drug-target datasets (i.e., activation and inhibition relationships given -1 and +1 labels, respectively) to modify the activity scores of pathways containing the target(s) of a drug. We describe the scoring algorithm in **Figure 2**.

**Figure 2.**
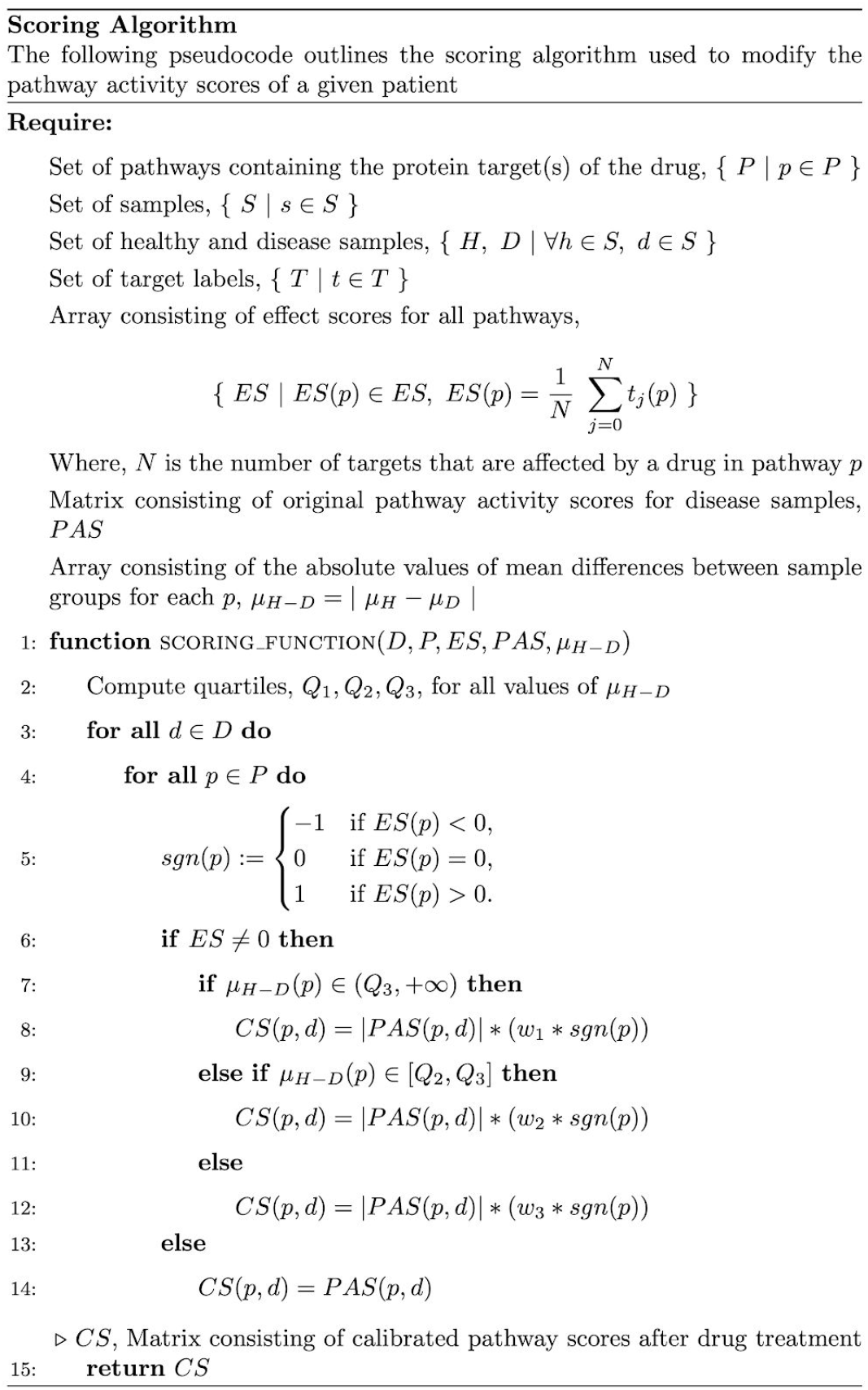
Scoring algorithm pseudocode. For each drug, if a given pathway contains one or more of its targets, the pathway is assigned an effect score *ES,* equivalent to the mean of the effect(s) (e.g., upregulated, downregulated or latent) of its target(s). The absolute values of the mean differences between healthy and disease groups are calculated for each pathway μ_H-D_*(p)* while their quartiles are then computed on line 2. Then, from lines 3-12, for each disease sample, if the *ES* of a pathway *p* is less than or greater than 0, the scoring algorithm calculates a calibration score *CS* as the product of the absolute value of the original pathway activity score *PAS,* the weight *w*, and the sign of the pathway *sgn(p).* We assign *w* based on the quartile μ_H-D_*(p)* pathway *p* falls into. For pathways with larger mean differences between groups, weights are assigned greater values, while pathways with smaller differences are weighted less. On lines 13-14, if the *ES* of a pathway *p* is 0, the *CS* is assigned the value of the original *PAS.* Finally, on line 15, the *CS* is returned as a score that simulates the effect of a drug on a pathway for a disease sample.

### 2.5. Drug response prediction and prioritization

The final step of our methodology aims at identifying drug candidates based on the predicted response of a patient to the simulated drug treatment. To do so, we input the modified features generated by the scoring algorithm in the trained ML model and re-evaluate the new class assignment of the patient.

Since the ML model has learnt to accurately differentiate between normal and disease samples, we expect that if a drug fails to affect a set of relevant pathways, the labels of the disease samples would remain unchanged. However, if the drug were to target a set of pathways dysregulated in a disease, we expect that the scoring algorithm could modify the scores so that they resemble those observed in control samples. Thus, by inputting these modified scores into the trained ML model, we can assess whether disease samples can now be classified as normal. Finally, after re-evaluating the predictions made by the ML model, we can rank promising drugs by the proportion of disease samples that are classified as normal as a proxy of the effectiveness of the drug.

### 2.6. Validation and robustness analysis

In this section, we outline the robustness experiments conducted to assess the ability of our methodology to identify drugs which have already been approved or tested in clinical trials for each of the three cancer types (i.e., TCGA datasets).

First, to simulate drug treatment using the scoring algorithm described in **Subsection 2.4**, we used two different drug-target datasets: DrugBank (Knox *et al.,* 2010) and DrugCentral (Ursu *et al*., 2016). For each of the datasets, we mapped drugs to DrugBank identifiers and protein targets to HGNC symbols. In total, we retrieved 1,346 unique drugs and 4,673 drug-target interactions from DrugBank and 638 unique drugs and 1,481 drug-target interactions from DrugCentral. We then used these drug-target interactions as the input to our methodology to simulate patient treatments **(Figure 1e)**.

For validation purposes, we used two ground-truth lists containing drug-disease pairs as true positives to verify the predictions made by our methodology **(Figure 1f)**. The first ground-truth list contained FDA-approved drugs for the three cancer types manually retrieved from the National Cancer Institute (https://www.cancer.gov/about-cancer/treatment/drugs/cancer-type) which we mapped to the two drug-target datasets previously described. The second ground-truth list contained drugs investigated in clinical trials for the three cancer datasets retrieved from the ClinicalTrials.gov website (downloaded on 16.04.2020). **Table 1** lists the number of approved and clinically tested drugs present in both drug-target datasets across the three investigated cancers.

**Table 1.** Number of FDA-approved and clinically tested drugs present in both drug-target datasets across the three investigated cancers. The percentage for the number of FDA-approved/clinically investigated drugs for each cancer type over the total number of drugs present in the drug-target dataset is reported between parentheses.

| - | Drug-target dataset |  |  |  |
| --- | --- | --- | --- | --- |
| - | DrugBank |  | DrugCentral |  |
| Dataset | Approved | Clinical trials | Approved | Clinical trials |
| BRCA | 26/1,346 (1.93%) | 182/1,346 (13.52%) | 14/638 (2.19%) | 115/638 (18.02%) |
| LIHC | 5/1,346 (0.37%) | 50/1,346 (3.71%) | 1/638 (0.16%) | 35/638 (5.49%) |
| PRAD | 13/1,346 (0.97%) | 134/1,346 (9.96%) | 7/638 (1.10%) | 84/638 (13.17%) |

As validation, both ground-truth lists were compared against the list of drugs that, according to our methodology, changed the predictions of 80% of the disease samples and subsequently classified them as normal. This threshold was selected as there were no drugs that changed the prediction for 90% or more of the patients with the parameters used by our scoring algorithm **(Supplementary Figures 3 and 4)**. We would like to note that the vast majority of the drugs will not change the predictions for most of the disease samples. Thus, we are exclusively interested in assessing the ability of our approach to recover true positives (i.e., positive predictive value) for the top-ranked drugs that change the predictions for a majority of patients. Furthermore, since our methodology aims to prioritize drug candidates, it suffers from an early retrieval problem (Berrar and Flach, 2012). Finally, only a small minority of drugs from the drug-target datasets can be used as positive labels for each of the indications, while the majority of drugs are not known to have therapeutic benefits for these indications; thus, creating a large imbalance between positive and negative labels. Due to these three reasons, the evaluation strategy we selected is more suitable than other conventional metrics such as the receiver operating characteristic (ROC) curves. Finally, to identify a set of weights for the three quartiles (i.e., *Q1, Q2 and Q3* (see **Figure 2**)) that perform well in the three datasets, we followed a similar strategy to Chen *et al*. (2016) where we tested different weight combinations with the intention of assigning larger weights to pathways with significantly higher dysregulations between cancer and normal samples (e.g., W_1_ > W_2_). By simultaneously conducting this parameter optimization on the three cancer datasets, we found a set of weights (i.e., *W_1_*=20, *W_2_*=5, and *W_3_*=10 for Q3 (the upper quartile representing the most dysregulated pathways), Q2 (middle quartile), and Q1 (lower quartile), respectively) that yielded a large proportion of true positives among the prioritized drugs and also performed better than any of the six methods we compared our methodology against below.

To test the robustness of our methodology, we replicated our experiments by generating one hundred sets of 1,346 drugs (the size of the DrugBank dataset) where each drug was assigned to a randomly selected protein target (from the set of all HGNC symbols) with a random causal effect (i.e., activation or inhibition). Next, we compared the number of drugs prioritized by these permutation experiments (i.e., drugs that changed the predictions of 80% of the disease samples) against the number of drugs prioritized by our methodology for the DrugBank dataset.

### 2.7. Performance comparison against equivalent drug-repurposing approaches

To evaluate our methodology, we compared it to six similar approaches that also employ transcriptomics data and pathway information to repurpose drugs on the BRCA and PRAD datasets (Chen *et al*., 2016; Chan *et al*. 2019) (note that the LIHC dataset is not included in their analyses). In the first of the two studies, Chan *et al.* (2019) evaluated the ability of their methodology and four additional approaches to predict known drugs (i.e., FDA-approved or in advanced clinical trials) for breast and prostate cancer. Similarly, Chen *et al.* (2016) reported the ability of their approach to identify FDA-approved drugs on the same datasets. We were thus able to directly compare the relative number of recovered true positives for each of these methods against ours.

### 2.8. Implementation and reproducibility

We performed GSEA with the Python package, GSEApy (version 0.9.12; https://github.com/zqfang/gseapy) and generated the ML models using scikit-learn (Pedregosa *et al*., 2011). The source code and data used in this manuscript is available at h ttps://github.com/sepehrgolriz/simdrugs under the Apache 2.0 License.

## 3. Results

### 3.1. Validation of the methodology and comparison against equivalent approaches

In this subsection, we investigate the drug candidates prioritized by our methodology in three different cancers and evaluate the ability of our approach to identify approved and clinically investigated drugs (i.e., true positives). **Table 2** shows that only a minority of the drugs present in both drug-target datasets were prioritized by our methodology given that a stringent threshold was employed which required that prioritized drugs change the predictions of at least 80% of the patients. Overall, our methodology is able to recover a large proportion of true positives (ranging from 15% to 32%) in all three cancers as well as in both drug-target datasets (**Table 2)**. This wide range may be attributable to a disproportion in the number of true positives that exist for each of the cancer datasets (e.g., BRCA has more than twice as many FDA-approved drugs and drugs in clinical trials than LIHC) as well as to the size of the drug-target datasets (i.e., DrugBank contains twice as many drugs as DrugCentral).

**Table 2.**
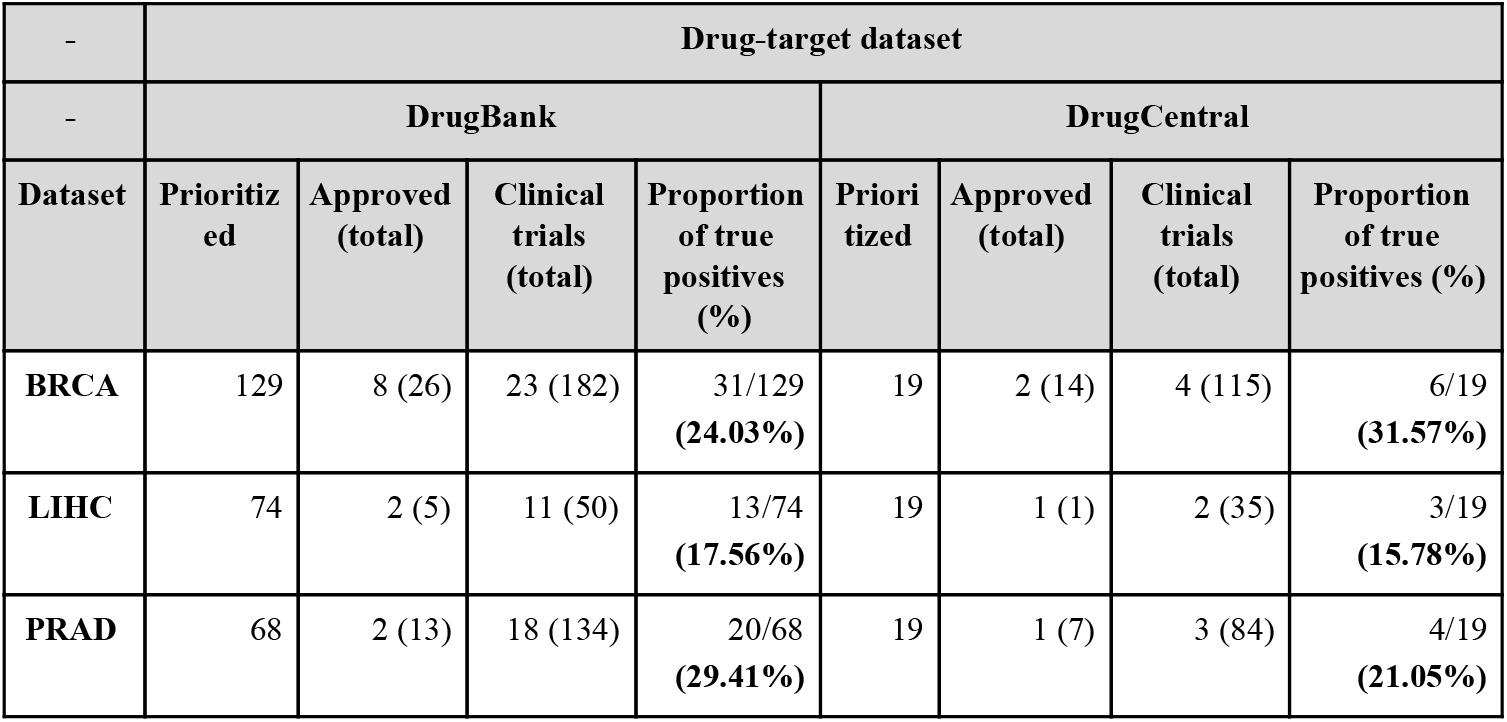
Number of FDA-approved and clinically tested drugs recovered for both drug-target datasets across the three investigated cancers. In the first column for each drug-target dataset (“Prioritized”), we report the number of drugs that changed the predictions for at least 80% of the patients for each cancer type. The second column (“Approved”) reports the number of **FDA**-approved drugs among these prioritized drugs as well as the total number of **FDA**-approved/clinically tested drugs present in each dataset between parentheses. Similarly, the third column (“Clinical trials”) reports the number of drugs tested in clinical trials among the prioritized drugs and the total number of **FDA**-approved/clinically tested drugs between parentheses. Finally, the last column (“Proportion of true positives”) reports the proportion of true positives (both **FDA**-approved and clinically tested drugs) among the prioritized drugs.

| - | Drug-target dataset |  |  |  |  |  |  |  |
| --- | --- | --- | --- | --- | --- | --- | --- | --- |
| - | DrugBank |  |  |  | DrugCentral |  |  |  |
| Dataset | Prioritized | Approved (total) | Clinical trials (total) | Proportion of true positives (%) | Prioritized | Approved (total) | Clinical trials (total) | Proportion of true positives (%) |
| BRCA | 129 | 8 (26) | 23 (182) | 31/129<br>(24.03%) | 19 | 2 (14) | 4 (115) | 6/19<br>(31.57%) |
| LIHC | 74 | 2 (5) | 11 (50) | 13/74<br>(17.56%) | 19 | 1 (1) | 2 (35) | 3/19<br>(15.78%) |
| PRAD | 68 | 2 (13) | 18 (134) | 20/68<br>(29.41%) | 19 | 1 (7) | 3 (84) | 4/19<br>(21.05%) |

As a comparison, the methodology proposed by Chan *et al*. (2019) reported lower proportions of true positives than our approach for the BRCA and PRAD datasets with 21.42% and 15.94%, respectively **(Supplementary Table 3)**. Furthermore, four additional methods present that were benchmarked by Chan *et al*. (2019) yielded even lower results on the same two cancer datasets **(Supplementary Tables 4-10)**. Similarly, Chen *et al*. (2016) also reported a lower proportion of true positive than our approach for the BRCA and PRAD datasets with 0.8% and 0.4%, respectively **(Supplementary Table 11).** Overall, the performance across all six methods varied from 0% to 11.53% for BRCA, and from 0.50% to 22.22% for PRAD and is summarized in **Supplementary Table 12**.

Additionally, the proportion of true positives yielded by our methodology is significantly higher than what one would expect by chance **(Table 1)**. Furthermore, we compared the number of prioritized drugs found in the original DrugBank and DrugCentral datasets to the number of prioritized drugs obtained in the robustness experiments in which we applied our methodology to drugs with randomly generated targets and target interactions **(Supplementary Figure 5)**. We found that all permutation experiments yielded a significantly lower number of prioritized drugs. Because our methodology can capture a much greater number of prioritized drugs on a real dataset, this validation highlights the capability of our approach to prioritize drugs with targets in relevant pathways that are key to change the predictions of patients.

### 3.2. In-depth investigation of the prioritized candidate drugs

Apart from the previous quantitative evaluation of our methodology, we conducted an in-depth analysis of the prioritized drugs to better understand the predictions made by our approach. Below, we focus on drugs prioritized using the DrugCentral dataset as this dataset contains a fewer number of prioritized drugs than DrugBank.

In the breast cancer dataset (BRCA), we identified a major class of drugs based on their mechanisms of action **(Figure 3a)**. This class targeted DNA and RNA metabolism and included commonly used anti-tumor drugs. One example of this group of drugs is fluorouracil, which targets thymidylate synthase, thereby, inhibiting the formation of thymidylate from uracil (Zhang *et al*., 2008). This drug is a chemotherapy medication commonly used to treat several cancers.

**Figure 3.**
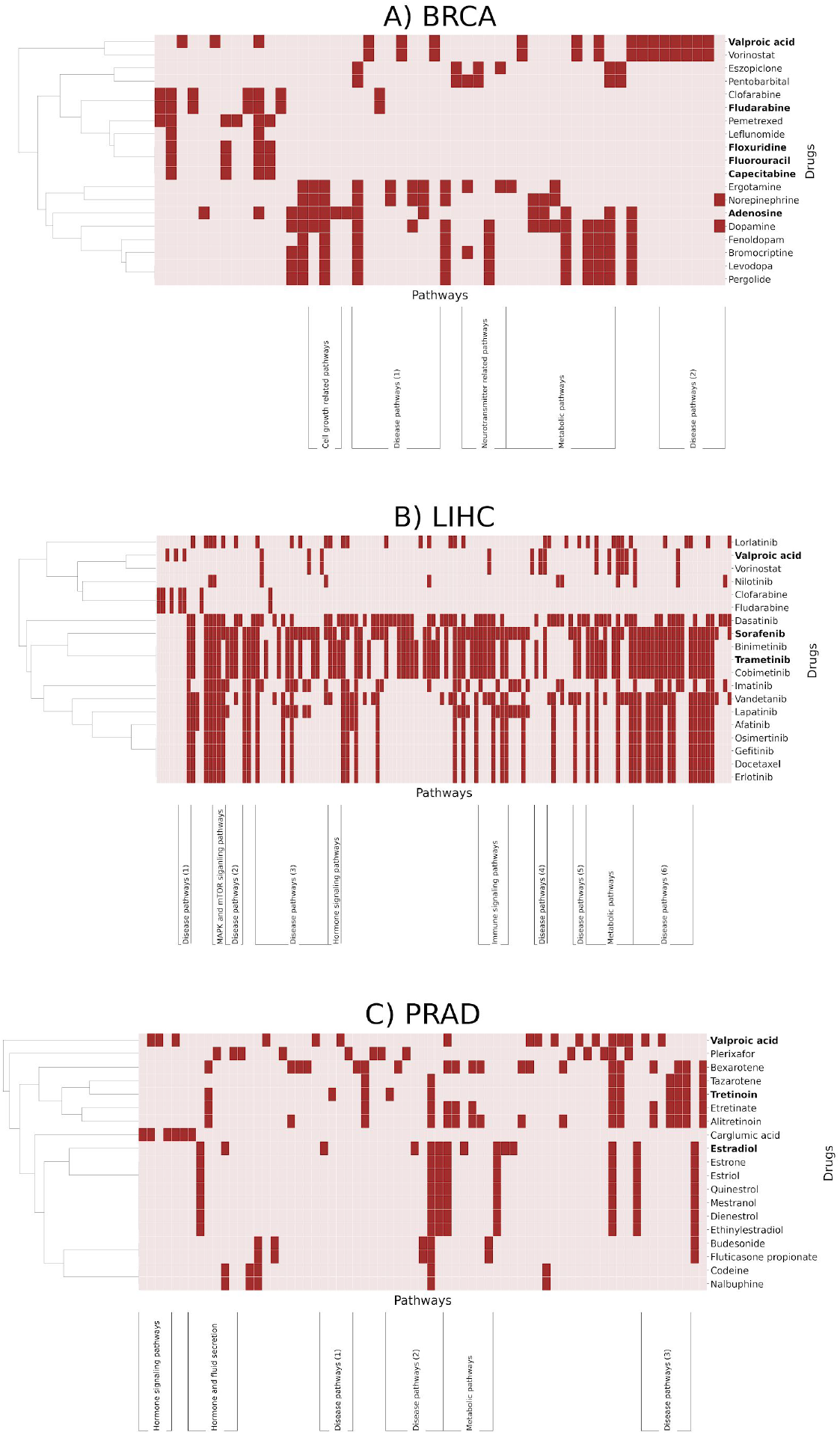
Pathways targeted by prioritized drugs in DrugCentral for each cancer dataset. The X-axis corresponds to pathways targeted by any of the prioritized drugs (i.e., pathways not targeted by any prioritized drug are omitted for better visualization). Prioritized drugs for each cancer dataset have been clustered based on the pathways they target and are reported on the Y-axis. Of the prioritized drugs, those that correspond to true positives are highlighted in bold. If a set of three or more similar pathways was clustered together, we manually assigned these pathways into distinct classes (Y-axis) Pathway names and cluster information are available as a **Supplementary File** and the equivalent figures for DrugBank are available as **Supplementary Figures 6-8.**

In the prostate cancer dataset (PRAD), we found that the majority of drugs were related to hormone metabolism and regulation **(Figure 3c)**. Due to the key role of sex steroid hormones in its initiation and progression (Snaterse *et al*., 2017), this cancer is classified as hormone-dependent. Thus, current treatments are often directly targeted towards these hormones, such as androgen deprivation therapy, which represents the major therapeutic option for treatment of advanced stages of this cancer (Harris *et al*., 2009; Karantanos *et al*., 2013).

The third dataset, LIHC, corresponds to hepatocarcinoma. Interestingly, the vast majority of the candidate drugs in this dataset (14/19) are tyrosine kinase inhibitors (TKI) corresponding to anti-tumor drugs already FDA-approved for other cancers (Huynh, 2010) **(Figure 3b)**. Since these kinases act as regulatory players in several cancer signaling pathways that can be hyperactivated, TKIs are used to “switch-off” these pathways, indirectly inhibiting cell growth (Khoo *et al*., 2019). One of the predicted drugs is sorafenib, which was the first TKI to be approved for the treatment of liver carcinoma and still remains as a first-line therapy. Similarly, another predicted drug, trametinib, is a dual-kinase inhibitor that is used in the treatment of advanced liver cancer. Finally, two of the remaining non-TKIs are also employed as chemotherapy drugs as they inhibit the synthesis of nucleotides.

### 3.3. Investigation of the prioritized cancer pathways

An additional application of our methodology is in pinpointing the pathway targets that are likely to be dysregulated in a disease. By investigating the drugs prioritized by our approach, using information on drug-target interactions and exploiting information revealed through the scoring algorithm, we can easily identify pathways that are targeted by the top-ranked drugs. We identified clusters of pathways belonging to several distinct classes **(Figure 3)**. We found that various metabolic pathways appear in all three datasets as the regulation of metabolism plays an important role in numerous cancers. Given that each of the three datasets were cancer subtypes, not surprisingly, we observed several disease-relevant pathways targeted by top-ranked drugs, among them approximately 30 KEGG cancer-related pathways (e.g., prostate cancer, pancreatic cancer, bladder cancer, and breast cancer).

Accordingly, we clustered the prioritized drugs from Figure 3 based on the pathways they target to assess whether drugs that target the same pathway are within the same class of drugs. Prioritized drugs for liver cancer can be clustered into four different classes of tyrosine kinase inhibitors: i) JAK inhibitors (i.e., sorafenib, vandetanib, erlotinib, and lapatinib), ii) ALK inhibitors (i.e., lorlatinib), iii) BCR–Abl (i.e., nilotinib, dasatinib, and imatinib), and iv) and EGFR inhibitors (i.e., afatinib) (Bhullar *et al.,* 2018). Additionally, we found MEK kinase inhibitors, specifically trametinib and cobimetinib. Finally, we found that while some drugs can change the predictions by targeting only a limited number of pathways (e.g., fludarabine in breast cancer and liver cancer), other drugs can change predictions by targeting several pathways (e.g., tretinoin in prostate cancer and trametinib in liver cancer).

Among the most commonly targeted pathways by the prioritized drugs in liver carcinoma, we found Ras/Raf/MAPK and PI3K/AKT/mTOR signaling, both of which have been reported to play important roles in the development of this type of cancer (Gedaly *et al.,* 2010). One of the prioritized drugs, sorafenib, is a multi-kinase inhibitor that targets several kinases including RFA1, PDGFR, and FLT3, which are involved in both tumor proliferation and angiogenesis (Mousa, 2008; Zhu *et al*., 2017). Sorafenib has been shown to inhibit tumor cell proliferation by blocking the Ras/Raf/MAPK pathway and to inhibit angiogenesis by blocking PDGFR signaling (Llovet *et al*., 2008) **(Supplementary Table 2)**.

### 3.4. Prioritizing combination therapies

Combination therapies are widely used for treating indications like cancer as they can often lead to the inhibition of the compensatory signaling pathways that maintain the growth and survival of tumor cells. In this section, we demonstrate how our methodology can be extended to predict the effects of a combination of drugs. To reduce the computational complexity associated with running our methodology on all possible combinations of drug pairs from both drug-target datasets (i.e., DrugBank and DrugCentral), we exclusively applied our method on all possible pairs from the set of prioritized drugs.

For two of the three datasets (i.e, LIHC and PRAD), nearly all drug pairs yielded better results (i.e., larger proportion of disease samples predicted as normal) than the use of a single drug alone. In the BRCA dataset, however, multiple combinations yielded worse results than those observed with single drug therapy. For example, the combination of bromocriptine with valproic acid decreased the proportion of treated patients from 80% to less than 10%. Specifically, bromocriptine is an adrenergic receptor agonist that stimulates the beta-adrenergic signaling pathway, which in turn prompts tumor angiogenesis and cancer development (Chen *et al.*, 2014). Similarly, valproic acid is a histone deacetylase which also induces beta-adrenergic signaling, thus promoting cancer progression (Hulsurkar *et al.*, 2017). Therefore, the combination of these two drugs not only fails to treat the cancer, but may in fact lead to the worsening of the condition.

## 4. Discussion

Here, we have presented a powerful machine learning framework to simulate drug responses for applications in drug discovery, precision medicine, and target identification. We demonstrate our methodology on three different cancer datasets by using patient-specific pathway signatures to train highly predictive models which we use as a proxy for drug candidate identification. Our results yielded a larger proportion of FDA-approved drugs as well as drugs investigated in clinical trials than six comparable approaches for the indications we studied, suggesting that other drugs prioritized by our methodology may also represent promising candidates for repurposing. Additionally,in contrast to the other methodologies, our approach is able to prioritize drugs for individual patients, making it suitable for precision medicine applications. Finally, we also show how our methodology can be applied to propose drug combinations as well as to reveal sets of dysregulated pathways that could be used as possible targets.

Currently, there exist several limitations to this study; first, although our scoring algorithm used to simulate drug response has been shown to yield promising results in the three datasets analyzed, other scoring algorithms may be better suited for different datasets and/or applications. For instance, we could tailor the current scoring algorithm for drug discovery to learn pathway signatures from approved drugs and use these drugs to prioritize candidates that exhibit similar patterns of activity. Second, although we used the same weights for all experiments on each of the three datasets, it may be the case that weights must be tuned for other datasets to yield promising candidates. Third, since our methodology relies on pathway signatures derived from transcriptomics data, it is inherently limited to indications where this modality is highly predictive. Thus, it would be less effective in indications where transcriptomics have limited prediction power to discriminate between normal and disease samples, such as Parkinson’s disease (Chen-Plotkin, 2018).

Beyond this proof-of-concept, our methodology can be extended to include several additional functionalities. For instance, drug administration could be simulated in an ML model that takes into consideration temporal dimensions (e.g., event-based models (Fonteijn H. M., *et al*. (2012), survival analysis (Tibshirani, 1997)). Furthermore, in this paper we trained a simple ML model, nonetheless, the same strategy could be applied to more complex ML or AI models. Since the elastic net penalty encourages sparsity, one may also use the coefficients of an ML model as a preliminary method of filtering for significant features. To save time, the total set of drug candidates can be subset to only those which directly affect the features that significantly affect the prediction capabilities of the model. Additionally, we restricted our analysis to a single pathway database as it was sufficient to deploy a predictive ML model for the specific classification task we presented. However, by incorporating pathway information from other databases into the ML model, we can increase the total number and coverage of pathways to potentially reveal additional pathway targets. Similarly, the use of different drug-target databases such as ExCAPE-DB (Sun *et al*., 2017) could broaden the chemical space and lead to the identification of new candidates. By combining brute-force and reverse engineering approaches, one can also identify the most effective pathway scores a drug should target for any given indication; thus, tailoring the presented methodology towards drug discovery. Finally, our methodology could also be deployed to support clinical decision-making in personalized medicine by simulating the effect of drugs on individual patients.

## Authors’ Contributions

DDF conceived and designed the study. SGK implemented the scoring algorithm and ran the validation experiments with assistance from DDF. SGK analyzed the case scenario and MHA and DDF assisted in the interpretation of the results. SM processed and prepared the datasets. SM and SGK ran the datasets with the pathway enrichment method to generate the pathway activity scores. SM and VSB trained the ML models. ATK, MHA, and DDF acquired the funding. SGK, SM, and DDF wrote the manuscript.

All authors have read and approved the final manuscript.

## Acknowledgements

The authors would like to thank Colin Birkenbihl and Lauren DeLong for their valuable feedback.

## References

1. Adam G., et al. (2020). Machine learning approaches to drug response prediction: challenges and recent progress. npj precision oncology, 4(1), 1–10. https://doi.org/10.1038/s41698-020-0122-1

2. Amadoz A., et al. (2019). A comparison of mechanistic signaling pathway activity analysis methods. Briefings in bioinformatics, 20(5), 1655–1668. https://doi.org/10.1093/bib/bby040

3. Ammad-ud-din M., et al. (2016). Drug response prediction by inferring pathway-response associations with kernelized Bayesian matrix factorization. Bioinformatics, 32(17), i455–i463. https://doi.org/10.1093/bioinformatics/btw433

4. Barbie D. A., et al. (2009). Systematic RNA interference reveals that oncogenic KRAS-driven cancers require TBK1. Nature, 462(7269), 108.https://doiorg/10.1038/nature08460

5. Berrar D., and Flach P. (2012). Caveats and pitfalls of ROC analysis in clinical microarray research (and how to avoid them). Briefings in bioinformatics, 13(1), 83–97. https://doi.org/10.1093/bib/bbr008

6. Bhullar K. S., et al. (2018). Kinase-targeted cancer therapies: progress, challenges and future directions. Molecular cancer, 17(1), 1–20. https://doi.org/10.1186/s12943-018-0804-2

7. Castro, V. M., et al. (2017). Large-scale identification of patients with cerebral aneurysms using natural language processing. Neurology, 88(2), 164–168. https://doi.org/10.1212/WNL.0000000000003490

8. Chan J., Wang X., Turner, J. A., Baldwin N. E., and Gu J. (2019). Breaking the paradigm: Dr Insight empowers signature-free, enhanced drug repurposing. Bioinformatics, 35(16), 2818–2826. https://doi.org/10.1093/bioinformatics/btz006

9. Chen H. R., Sherr D. H., Hu Z., and DeLisi, C. (2016). A network based approach to drug repositioning identifies plausible candidates for breast cancer and prostate cancer. BMC medical genomics, 9(1), 1–11. https://doi.org/10.1186/s12920-016-0212-7

10. Chen-Plotkin A. S. (2018). Blood transcriptomics for Parkinson disease?. Nature Reviews Neurology, 14(1), 5–6. https://doi.org/10.1038/nrneurol.2017.166

11. Chen H., et al. (2014). Adrenergic signaling promotes angiogenesis through endothelial cell-tumor cell crosstalk. Endocrine-Related Cancer, 21(5), 783–95. https://doi.org/10.1530/erc-14-0236

12. Colaprico A., et al. (2015). TCGAbiolinks: an R/Bioconductor package for integrative analysis of TCGA data. Nucleic acids research 44(8), e71–e71. https://doi.org/10.1093/nar/gkv1507

13. Emon M. A., et al. (2020). PS4DR: A multimodal workflow for identification and prioritization of drugs based on pathway signatures. BMC bioinformatics, 21(1), 1–21. https://doi.org/10.1186/s12859-020-03568-5

14. Esteban-Medina M., et al. (2019). Exploring the druggable space around the Fanconi anemia pathway using machine learning and mechanistic models. BMC bioinformatics, 20(1), 370. https://doi.org/10.1186/s12859-019-2969-0

15. Fonteijn H. M., et al. (2012). An event-based model for disease progression and its application in familial Alzheimer’s disease and Huntington’s disease. NeuroImage, 60(3), 1880-1889 https://doi.org/10.1007/978-3-642-22092-0_61

16. Gedaly R., et al. (2010). PI-103 and sorafenib inhibit hepatocellular carcinoma cell proliferation by blocking Ras/Raf/MAPK and PI3K/AKT/mTOR pathways. Anticancer research, 30(12), 4951–4958.

17. Harris W. P., et al. (2009). Androgen deprivation therapy: progress in understanding mechanisms of resistance and optimizing androgen depletion. Nature clinical practice Urology, 6(2), 76–85. https://doiorg/10.1038/ncpuro1296

18. Hirohara M., et al. (2018). Convolutional neural network based on SMILES representation of compounds for detecting chemical motifs. BMC bioinformatics, 19(19), 526. https://doi.org/10.1186/s12859-018-2523-5

19. Hulsurkar M., et al. (2017). Beta-adrenergic signaling promotes tumor angiogenesis and prostate cancer progression through HDAC2-mediated suppression of thrombospondin-1. Oncogene, 36(11), 1525–1536. https://doi.org/10.1038/onc.2016.319

20. Huynh H. (2010). Tyrosine kinase inhibitors to treat liver cancer. Expert opinion on emerging drugs, 15(1), 13–26. https://doi.org/10.1517/14728210903571659

21. Jassal B., et al. (2020). The reactome pathway knowledgebase. Nucleic acids research, 48(D1), D498–D503. https://doi.org/10.1093/nar/gkz1031

22. Kanehisa M., Furumichi M., Tanabe M., Sato Y., and Morishima K. (2017). KEGG: new perspectives on genomes, pathways, diseases and drugs. Nucleic acids research, 45(D1), D353–D361. https://doi.org/10.1093/nar/gkw1092

23. Karantanos T., Corn P. G., and Thompson T. C. (2013). Prostate cancer progression after androgen deprivation therapy: mechanisms of castrate resistance and novel therapeutic approaches. Oncogene, 32(49), 5501–5511. https://doi.org/10.1038/onc.2013.206

24. Khoo T. S. W. L., Rehman A., and Olynyk J. K. (2019). Tyrosine Kinase Inhibitors in the Treatment of Hepatocellular Carcinoma. Exon Publications, 127–139.

25. Knox C., et al. (2010). DrugBank 3.0: a comprehensive resource for ‘omics’ research on drugs. Nucleic acids research, 39(suppl_1), D1035–D1041. https://doi.org/10.1093/nar/gkq1126

26. Lim S., et al. (2020). Comprehensive and critical evaluation of individualized pathway activity measurement tools on pan-cancer data. Briefings in bioinformatics, 21(1), 36–46. https://doi.org/10.1093/bib/bby097

27. Liu S., et al. (2014). Early diagnosis of Alzheimer’s disease with deep learning. IEEE 11th international symposium on biomedical imaging (ISBI), 1015–1018). https://doiorg/10.1109/ISBI.2014.6868045

28. Llovet J. M., et al. (2008). Sorafenib in advanced hepatocellular carcinoma. New England journal of medicine, 359(4), 378–390. https://doi.org/10.1056/nejmoa0708857

29. Molinaro A. M., Simon R., and Pfeiffer R. M. (2005). Prediction error estimation: a comparison of resampling methods. Bioinformatics, 21(15), 3301–3307. https://doi.org/10.1093/bioinformatics/bti499

30. Mousa A. B. (2008). Sorafenib in the treatment of advanced hepatocellular carcinoma. Saudi journal of gastroenterology: official journal of the Saudi Gastroenterology Association, 14(1), 40. https://doi.org/10.4103/1319-3767.37808

31. Mubeen S., et al. (2019). The impact of pathway database choice on statistical enrichment analysis and predictive modeling. Frontiers in genetics, 10, 1203. https://doi.org/10.3389/fgene.2019.01203

32. Nguyen, T. M., Shafi, A., Nguyen, T., and Draghici, S. (2019). Identifying significantly impacted pathways: a comprehensive review and assessment. Genome biology, 20(1), 1–15. https://doi.org/10.1186/s13059-019-1790-4

33. Pai S., et al. (2019). netDx: Interpretable patient classification using integrated patient similarity networks. Molecular systems biology, 15(3). https://doi.org/10.15252/msb.20188497

34. Pedregosa F., et al. (2011). Scikit-learn: Machine learning in Python. The Journal of machine Learning research, 12, 2825–2830.

35. Peyvandipour A., Saberian N., Shafi A., Donato M., and Draghici, S. (2018). A novel computational approach for drug repurposing using systems biology. Bioinformatics, 34(16), 2817–2825. https://doi.org/10.1093/bioinformatics/bty133

36. Perscheid C. (2020). Integrative biomarker detection on high-dimensional gene expression data sets: a survey on prior knowledge approaches. Briefings in bioinformatics, bbaa151. https://doi.org/10.1093/bib/bbaa151

37. Réda C., et al. (2020). Machine learning applications in drug development. Computational and Structural Biotechnology Journal, 18, 241–252. https://doi.org/10.1016/j.csbj.2019.12.006

38. Reimand J., et al. (2019). Pathway enrichment analysis and visualization of omics data using g: Profiler, GSEA, Cytoscape and EnrichmentMap. Nature protocols, 14, 482–517. https://doi.org/10.1038/s41596-018-0103-9

39. Saberian N., Peyvandipour A., Donato M., Ansari S., and Draghici, S. (2019). A new computational drug repurposing method using established disease–drug pair knowledge. Bioinformatics, 35(19), 3672–3678. https://doi.org/10.1093/bioinformatics/btz156

40. Snaterse G., et al. (2017). Circulating steroid hormone variations throughout different stages of prostate cancer. Endocrine-Related Cancer, 24(11), R403–R420. https://doiorg/10.1530/erc-17-0155

41. Su J., Yoon B. J., and Dougherty E. R. (2009). Accurate and reliable cancer classification based on probabilistic inference of pathway activity. PloS one, 4(12), e8161. https://doi.org/10.1371/journal.pone.0008161

42. Sun J., et al. (2017). ExCAPE-DB: an integrated large scale dataset facilitating Big Data analysis in chemogenomics. Journal of cheminformatics, 9(1), 17. https://doi.org/10.1186/s13321-017-0203-5

43. Tibshirani R. (1997). The lasso method for variable selection in the Cox model. Statistics in medicine, 16(4), 385–395.

44. Ursu O., et al. (2016). DrugCentral: online drug compendium. Nucleic acids research, 45(D1), D932–D939. https://doi.org/10.1093/nar/gkw993.

45. Wang X., et al. (2019). Predict drug sensitivity of cancer cells with pathway activity inference. BMC medical genomics, 12(1), 5–13. https://doi.org/10.1186/s12920-018-0449-4

46. Weinstein J. N., et al. (2013). The cancer genome atlas pan-cancer analysis project. Nature genetics, 45(10), 1113. https://doi.org/10.1038/ng.2764

47. Zhang N., et al. (2008). 5-Fluorouracil: mechanisms of resistance and reversal strategies. Molecules, 13(8), 1551–1569. https://doi.org/10.3390/molecules13081551

48. Zhao K. and So H. C. (2019). Using drug expression profiles and machine learning approach for drug repurposing. Computational methods for drug repurposing, 219–237. Humana Press, New York, NY. https://doi.org/10.1007/978-1-4939-8955-3_13

49. Zhu Y. J., Zheng B., Wang H. Y., and Chen L. (2017). New knowledge of the mechanisms of sorafenib resistance in liver cancer. Acta pharmacologica Sinica, 38(5), 614–622. https://doi.org/10.1038/aps.2017.5

