## Supplementary File 1 for "Using predictive machine learning models for drug response simulation by calibrating patient-specific pathway signatures"

#### Supplementary Tables

| Dataset | Normal samples | Tumor samples |
|---|---|---|
| **BRCA** | 113 | 1102 |
| **LIHC** | 50 | 371 |
| **PRAD** | 52 | 498 |

**Supplementary Table 1. Number of normal and tumor samples in the TCGA datasets used in this work.**

| Pathway | Target | Pathway- level drug effect | Pathway activity in patients relative to controls |
|---|---|---|---|
| Gap junction | PDGFRB, RAF1 | Inhibition | Upregulated |
| Fc gamma R mediated phagocytosis | RAF1 | Inhibition | Upregulated |
| Phospholipase D signaling pathway | INSR, PDGFRB, KIT, RAF1 | Inhibition | Upregulated |
| Thyroid hormone signaling pathway | RAF1 | Inhibition | Upregulated |
| Thyroid cancer | BRAF, RET | Inhibition | Upregulated |
| Hepatitis B | BRAF, RAF1 | Inhibition | Upregulated |
| Human papillomavirus infection | PDGFRB, RAF1 | Inhibition | Upregulated |
| Focal adhesion | BRAF, FLT4, KDR, PDGFRB, RAF1, FLT1 | Inhibition | Upregulated |
| Renal cell carcinoma | BRAF, RAF1 | Inhibition | Upregulated |
| Glioma | BRAF, PDGFRB, RAF1 | Inhibition | Upregulated |
| Axon guidance | RAF1 | Inhibition | Upregulated |
| Endocrine resistance | BRAF, RAF1 | Inhibition | Upregulated |
| Breast cancer | BRAF, FLT4, KIT, RAF1 | Inhibition | Upregulated |
| Sphingolipid signaling pathway | RAF1 | Inhibition | Upregulated |
| Autophagy animal | RAF1 | Inhibition | Upregulated |
| mTOR signaling pathway | INSR, BRAF, RAF1 | Inhibition | Upregulated |
| GnRH signaling pathway | RAF1 | Inhibition | Upregulated |
| Pathways in cancer | FLT3, BRAF, FLT4, PDGFRB, KIT, RAF1, RET | Inhibition | Upregulated |
| Chronic myeloid leukemia | BRAF, RAF1 | Inhibition | Upregulated |
| Choline metabolism in cancer | PDGFRB, RAF1 | Inhibition | Upregulated |
| Bladder cancer | BRAF, RAF1 | Inhibition | Upregulated |
| Non small cell lung cancer | BRAF, RAF1 | Inhibition | Upregulated |
| Gastric cancer | BRAF, RAF1 | Inhibition | Upregulated |
| Cushing syndrome | BRAF | Inhibition | Upregulated |
| VEGF signaling pathway | KDR, RAF1 | Inhibition | Upregulated |

| | | | |
|---|---|---|---|
| Hepatocellular carcinoma | BRAF, RAF1 | Inhibition | Upregulated |
| MAPK signaling pathway | INSR,FLT3,BRAF,FLT4,KDR,PDGFRB,KIT,RAF1, | Inhibition | Upregulated |
| Regulation of actin cytoskeleton | BRAF, PDGFRB, RAF1 | Inhibition | Upregulated |
| Human immunodeficiency virus 1 infection | RAF1 | Inhibition | Upregulated |
| Relaxin signaling pathway | RAF1 | Inhibition | Upregulated |
| Estrogen signaling pathway | RAF1 | Inhibition | Upregulated |
| Progesterone mediated oocyte maturation | BRAF, RAF1 | Inhibition | Upregulated |
| MicroRNAs in cancer | PDGFRB, RAF1 | Inhibition | Upregulated |
| Neurotrophin signaling pathway | BRAF, RAF1 | Inhibition | Upregulated |
| Alcoholism | BRAF, RAF1 | Inhibition | Upregulated |
| Fc epsilon RI signaling pathway | RAF1 | Inhibition | Upregulated |
| Apoptosis | RAF1 | Inhibition | Upregulated |
| Cellular senescence | RAF1 | Inhibition | Upregulated |
| Colorectal cancer | BRAF, RAF1 | Inhibition | Upregulated |
| Long term depression | BRAF, RAF1 | Inhibition | Upregulated |
| Melanogenesis | KIT, RAF1 | Inhibition | Upregulated |

**Supplementary Table 2. Effect of Sorafenib on pathway targets in the LIHC dataset.** The first column corresponds to the pathways that contain protein targets of Sorafenib while the second column corresponds to the specific protein targets of the drug. The third column corresponds to the effect of the drug on the pathway based on its effect on the target. In this case, all pathways are inhibited as all protein targets are inhibited by Sorafenib. Finally, the last column presents a relative comparison between the pathway activity observed in patients vs. controls: downregulated corresponds to lower pathway activity and upregulated corresponds to the opposite.

| **Dr. Insight Performance** | | |
|---|---|---|
| **Dataset** | **TCGA BRCA** | **TCGA PRAD** |
| **# identified drug treatments** | 70 | 69 |
| **# identified ground-truth drug treatments** | 15 | 11 |
| **Proportion of true positives (%)** | **21.42** | **15.94** |

**Supplementary Table 3. Number of approved drugs recovered using Dr. Insight on the BRCA and PRAD dataset.** The results of this table are reported in Chan *et al*. (2019) (Table S5.11) https://doi.org/10.1093/bioinformatics/btz006.

| **CMap performance on BRCA** | | | | | | | | |
|---|---|---|---|---|---|---|---|---|
| **Gene signature sizes (threshold)** | 50 | 100 | 200 | 300 | 400 | 600 | 800 | 1000 |
| **# identified drug treatments** | 56 | 52 | 55 | 67 | 74 | 74 | 71 | 56 |
| **# identified ground-truth drug treatments** | 5 | 6 | 3 | 6 | 5 | 6 | 6 | 5 |
| **Proportion of true positives (%)** | 8.92 | **11.53** | 5.45 | 8.92 | 6.75 | 8.1 | 8.45 | 8.92 |

**Supplementary Table 4. Number of approved drugs recovered using CMap on the BRCA dataset.** The proportion of true positives highlighted in bold indicates the highest percentage amongst all parameters (i.e., threshold gene set size). The results of this table are reported by Chan *et al*. (2019) (Table S5.1) https://doi.org/10.1093/bioinformatics/btz006.

| CMap performance on PRAD | | | | | | | | |
|---|---|---|---|---|---|---|---|---|
| **Gene signature sizes (threshold)** | 50 | 100 | 200 | 300 | 400 | 600 | 800 | 1000 |
| **# identified drug treatments** | 52 | 83 | 65 | 66 | 80 | 73 | 76 | 71 |
| **# identified ground-truth drug treatments** | 8 | 11 | 7 | 7 | 9 | 10 | 8 | 9 |
| **Proportion of true positives (%)** | **15.38** | 13.25 | 10.76 | 10.60 | 11.25 | 13.69 | 10.52 | 12.67 |

**Supplementary Table 5. Number of approved drugs recovered using CMap on the PRAD dataset.** The proportion of true positives highlighted in bold indicates the highest percentage amongst all parameters (i.e., threshold gene set size). The results of this table are reported by Chan *et al.* (2019) (Table S5.2) https://doi.org/10.1093/bioinformatics/btz006.

| sscMap performance on BRCA | | | | | | | | |
|---|---|---|---|---|---|---|---|---|
| **Gene signature sizes (threshold)** | 50 | 100 | 200 | 300 | 400 | 600 | 800 | 1000 |
| **# identified drug treatments** | 6 | 19 | 659 | 998 | 1185 | 1316 | 1404 | 1436 |
| **# identified ground-truth drug treatments** | 0 | 1 | 31 | 45 | 61 | 63 | 72 | 77 |
| **Proportion of true positives (%)** | 0 | 5.26 | 4.70 | 4.50 | 5.14 | 4.78 | 5.12 | **5.36** |

**Supplementary Table 6. Number of approved drugs recovered using sscMap on the BRCA dataset.** The proportion of true positives highlighted in bold indicates the highest percentage amongst all parameters (i.e., threshold gene set size). The results of this table are reported by Chan *et al.* (2019) (Table S5.4) https://doi.org/10.1093/bioinformatics/btz006.

| sscMap performance on PRAD | | | | | | | | |
|---|---|---|---|---|---|---|---|---|
| **Gene signature sizes (threshold)** | 50 | 100 | 200 | 300 | 400 | 600 | 800 | 1000 |
| **# identified drug treatments** | 8 | 18 | 100 | 177 | 202 | 381 | 653 | 810 |
| **# identified ground-truth drug treatments** | 1 | 4 | 9 | 11 | 14 | 17 | 21 | 25 |
| **Proportion of true positives (%)** | 12.5 | **22.22** | 9 | 6.21 | 6.93 | 4.46 | 3.21 | 3.08 |

**Supplementary Table 7. Number of approved drugs recovered using sscMap on the BRCA dataset.** The proportion of true positives highlighted in bold indicates the highest percentage amongst all parameters (i.e., threshold gene set size). The results of this table are reported by Chan *et al.* (2019) (Table S5.5) https://doi.org/10.1093/bioinformatics/btz006.

| NFFinder performance on BRCA | | | | | | | | |
|---|---|---|---|---|---|---|---|---|
| **Gene signature sizes (threshold)** | 50 | 100 | 200 | 300 | 400 | 600 | 800 | 1000 |
| **# identified drug treatments** | 329 | 285 | 623 | 782 | 651 | 854 | 1007 | 1069 |
| **# identified ground-truth drug treatments** | 26 | 24 | 40 | 45 | 43 | 54 | 66 | 62 |
| **Proportion of true positives (%)** | 7.90 | **8.42** | 6.42 | 5.75 | 6.60 | 6.32 | 6.65 | 5.59 |

**Supplementary Table 8. Number of approved drugs recovered using NFFinder on the BRCA dataset.** The proportion of true positives highlighted in bold indicates the highest percentage amongst all parameters (i.e., threshold gene set size). The results of this table are reported by Chan *et al.* (2019) (Table S5.7) https://doi.org/10.1093/bioinformatics/btz006.

| NFFinder performance on PRAD | | | | | | | | |
|---|---|---|---|---|---|---|---|---|
| **Gene signature sizes (threshold)** | 50 | 100 | 200 | 300 | 400 | 600 | 800 | 1000 |
| **# identified drug treatments** | 347 | 592 | 548 | 717 | 529 | 719 | 842 | 783 |
| **# identified ground-truth drug treatments** | 18 | 28 | 32 | 34 | 32 | 38 | 42 | 38 |
| **Proportion of true positives (%)** | 5.18 | 4.72 | **5.83** | 4.50 | 4.74 | 5.28 | 4.98 | 4.85 |

**Supplementary Table 9. Number of approved drugs recovered using NFFinder on the PRAD dataset.** The proportion of true positives highlighted in bold indicates the highest percentage amongst all parameters (i.e., threshold gene set size). The results of this table are reported by Chan *et al*. (2019) (Table S5.8) https://doi.org/10.1093/bioinformatics/btz006.

| Cogena | | |
|---|---|---|
| **Cogena** | TCGA BRCA | TCGA PRAD |
| **# identified drug treatments** | 335 | 982 |
| **# identified ground-truth drug treatments** | 30 | 5 |
| **Proportion of true positives (%)** | **8.95** | **0.50** |

**Supplementary Table 10. Number of approved drugs recovered using Cogena on the BRCA and PRAD dataset.** The results of this table are reported by Chan *et al*. (2019) (Table S5.10) https://doi.org/10.1093/bioinformatics/btz006.

| Chen *et al.* approach performance | | | |
|---|---|---|---|
| **Dataset** | **Prioritized** | **Approved (total)** | **Proportion of true positives (%)** |
| **BRCA** | 2435 | 20 (20) | 20/2435 **(0.81%)** |
| **PRAD** | 2500 | 10 (11) | 10/2500 **(0.40%)** |

**Supplementary Table 11. Number of approved drugs recovered reported by Chen *et al.* (2016) on the BRCA and PRAD dataset.** The results from this table are reported on Table 1 of the paper (https://doi.org/10.1186/s12920-016-0212-7).

| Method | BRCA | PRAD |
|---|---|---|
| **Dr. Insight** | 21.42 (%) | 15.94 (%) |
| **CMap** | 6.75 - 11.53 (%) | 10.52 - 15.38 (%) |
| **sscMap** | 0 - 5.36 (%) | 3.21 - 22.22 (%) |
| **NFFinder** | 5.75 - 8.42 (%) | 4.5 - 5.83 (%) |
| **Cogena** | 8.95 (%) | 0.50 (%) |
| **Chen *et al.*** | 0.81 (%) | 0.40 (%) |

**Supplementary Table 12. Performance for all methods benchmarked on the BRCA and PRAD dataset.** The performance is measured as % of approved drugs recovered and is taken from Supplementary Tables 3-11.

### Supplementary Figures

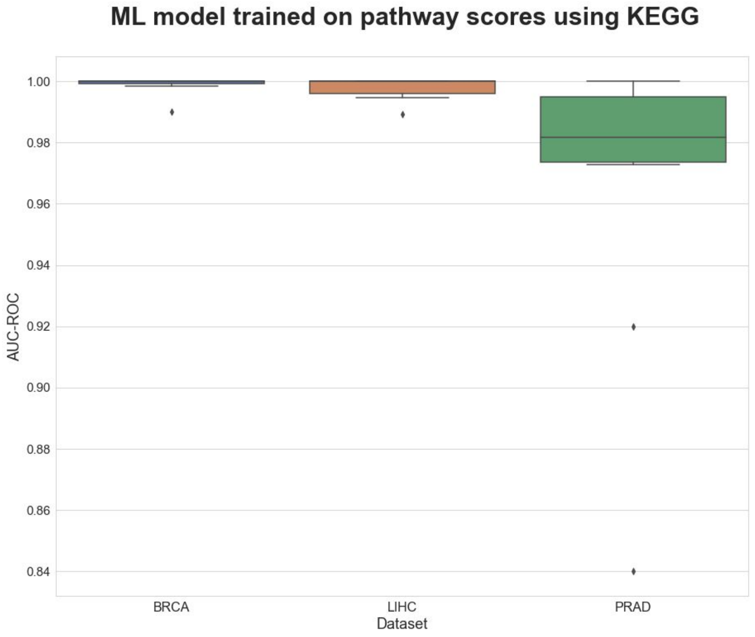


**Supplementary Figure 1. Prediction performance measured as AUC-ROC values of an elastic net classifier (tumor vs. normal samples) trained on the three TCGA datasets using pathway activity scores from ssGSEA run on KEGG.** Each boxplot shows the distribution of the AUCs over 10 repeats of the 10-fold cross-validation procedure. The same classifiers yielded equivalent AUC-PR values (data not shown).

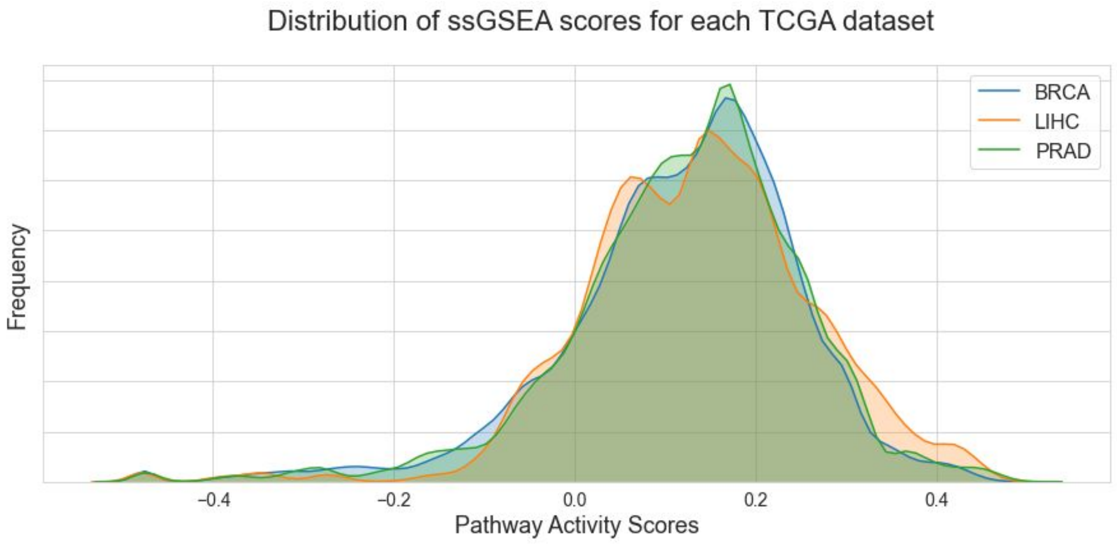


**Supplementary Figure 2. Distribution of pathway activity scores for each dataset.**

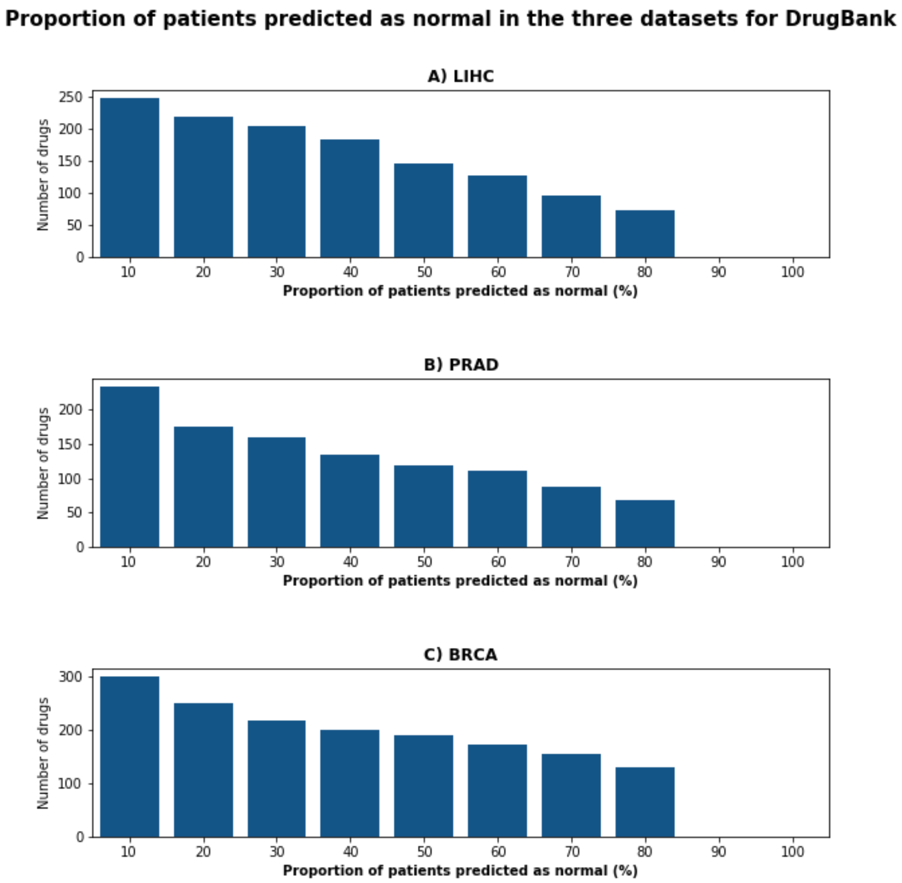


**Supplementary Figure 3. Proportion of patients predicted as normal for each cancer dataset using DrugBank.** Only a fraction of all drugs in DrugBank changed the predictions of 10% of the patients to normal. As the proportion of the samples changed increases, the number of prioritized drugs decreases to 19 for all three datasets.

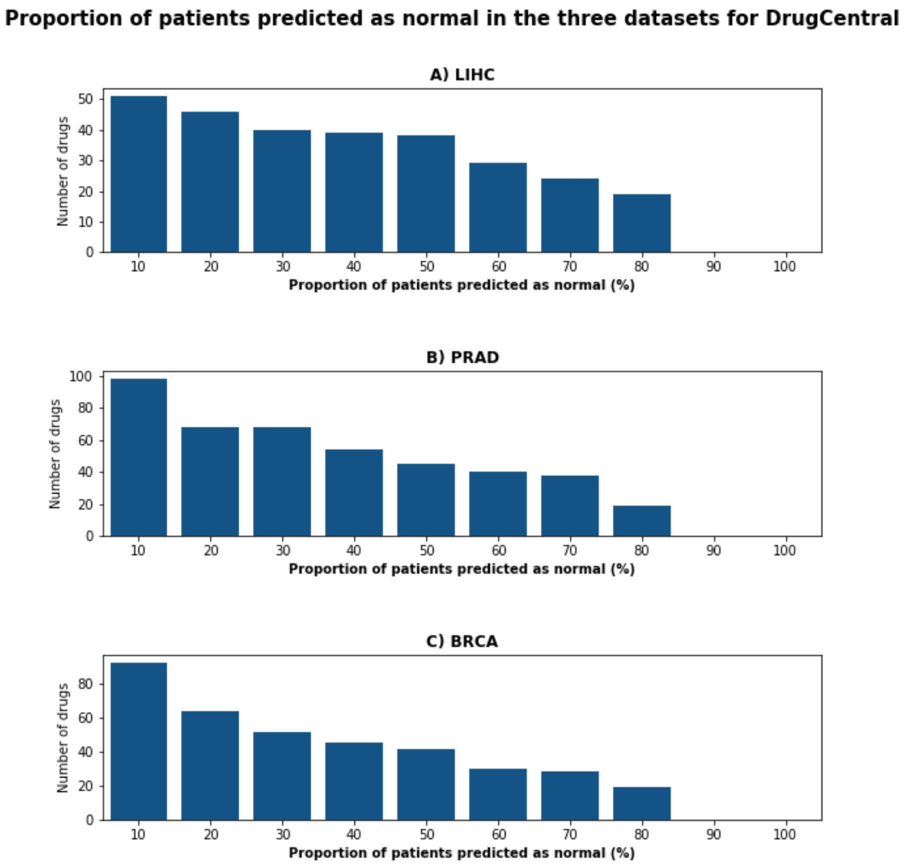


**Supplementary Figure 4. Proportion of patients predicted as normal for each cancer dataset using DrugCentral.** Only a fraction of all drugs in DrugCentral changed the predictions of 10% of the patients to normal. As the proportion of the samples changed increases, the number of prioritized drugs decreases to a shortlist of drugs for all three datasets.

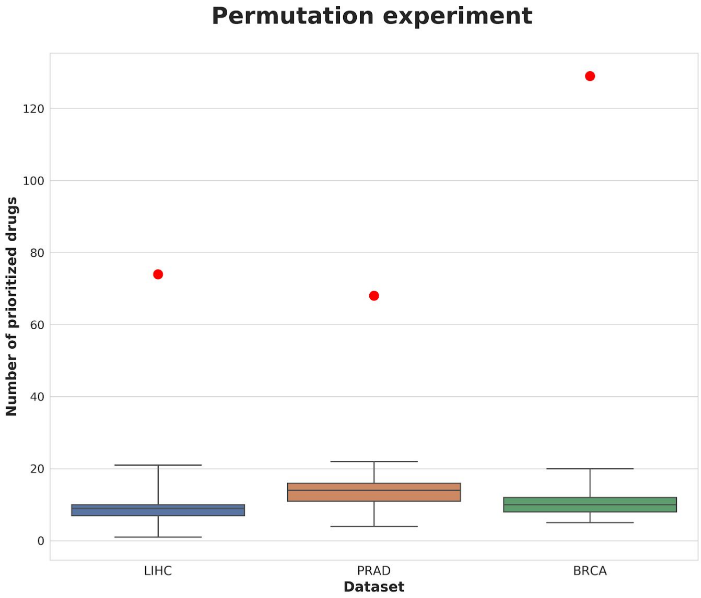


**Supplementary Figure 5. Comparison of the results of the permutation experiments against the number of prioritized drugs in DrugBank for the three cancer datasets.** While the three boxplots correspond to the number of prioritized drugs in the 100 permutation experiments for each of the three datasets, the number of prioritized drugs in the original DrugBank dataset has been indicated with a red circle. The number of prioritized drugs from DrugBank is significantly higher than for any of the permutations experiments. *p*-values have been omitted as all permutation experiments yielded a lower number of prioritized drugs compared to the original dataset and thus, *p*-values would be dominated by the number of experiments (i.e., 100 experiments would yield a *p*-value of 0.01, and 1,000 experiments would yield a *p*-value of 0.001). We would like to note that we compare the permutation experiments against DrugBank as the number of simulated drugs is equal to the size of this dataset (1,346 drugs). Furthermore, a comparison to the DrugCentral dataset would yield an even greater difference in the number of prioritized drugs as DrugCentral is smaller in size (638 drugs).

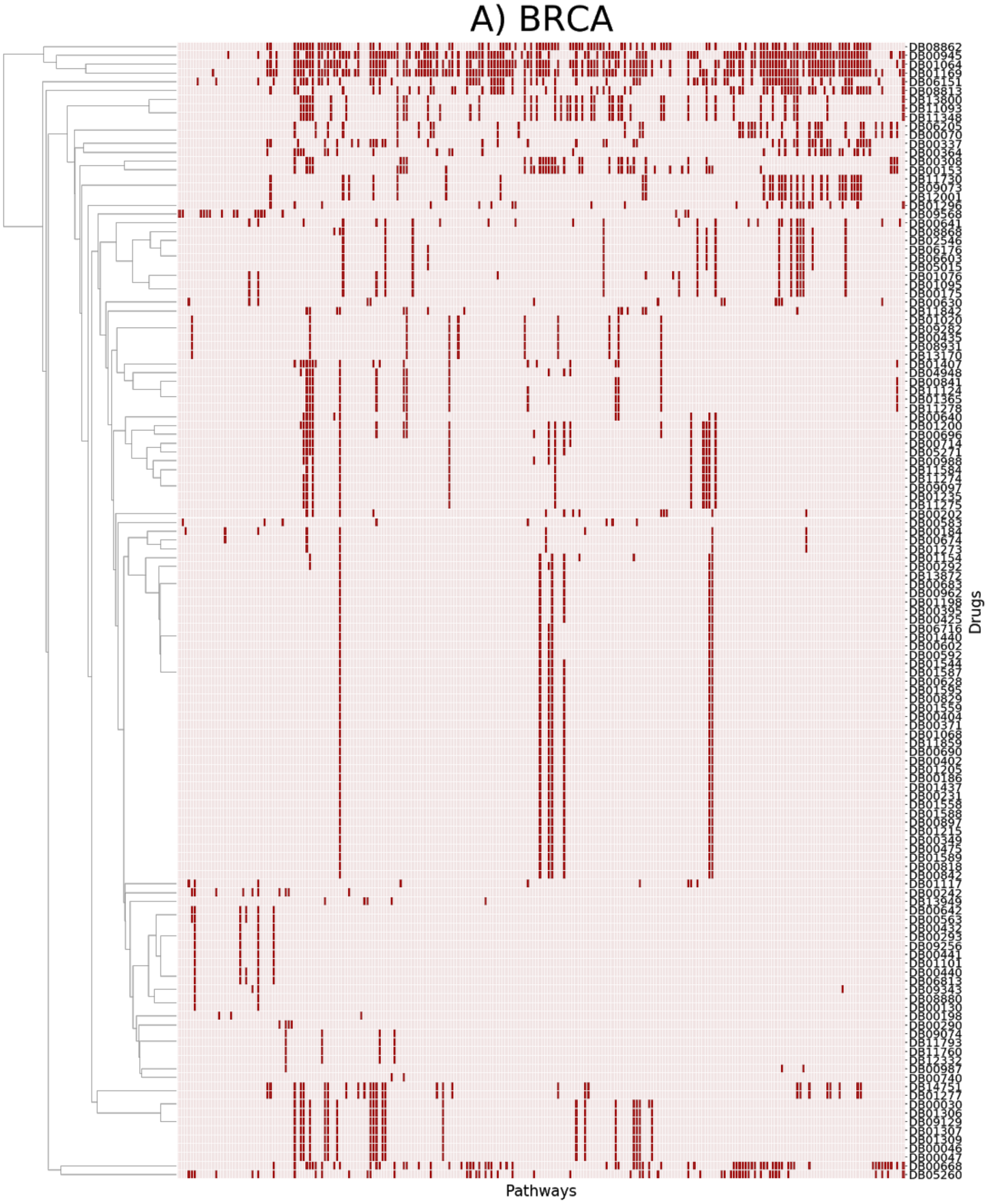


**Supplementary Figure 6. Pathways targeted by the prioritized drugs in DrugBank for BRCA.** The X-axis corresponds to the pathways targeted by any of the prioritized drugs (i.e., other KEGG pathways not targeted by any prioritized drug have been omitted for better visualization). Drugs (Y-axis) have been clustered based on the pathways they target. Due to the large number of pathways, we have clustered pathway groups together for visualization purposes. Black cells correspond to pathways targeted for each drug.

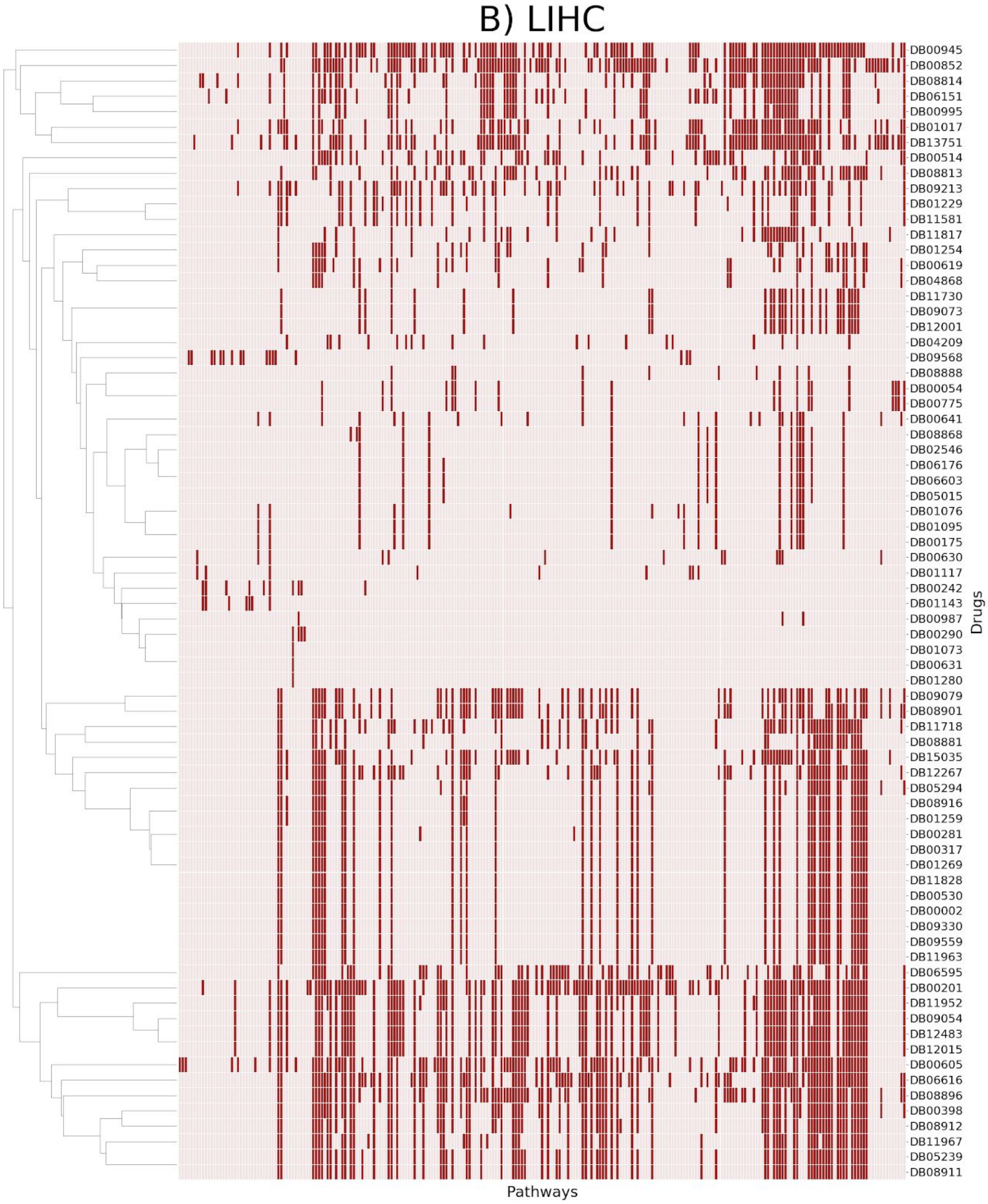


**Supplementary Figure 7. Pathways targeted by the prioritized drugs in DrugBank for LIHC.** The X-axis corresponds to the pathways targeted by any of the prioritized drugs (i.e., other KEGG pathways not targeted by any prioritized drug have been omitted for better visualization). Drugs (Y-axis) have been clustered based on the pathways they target. Due to the large number of pathways, we have clustered pathway groups together for visualization purposes. Black cells correspond to pathways targeted for each drug.

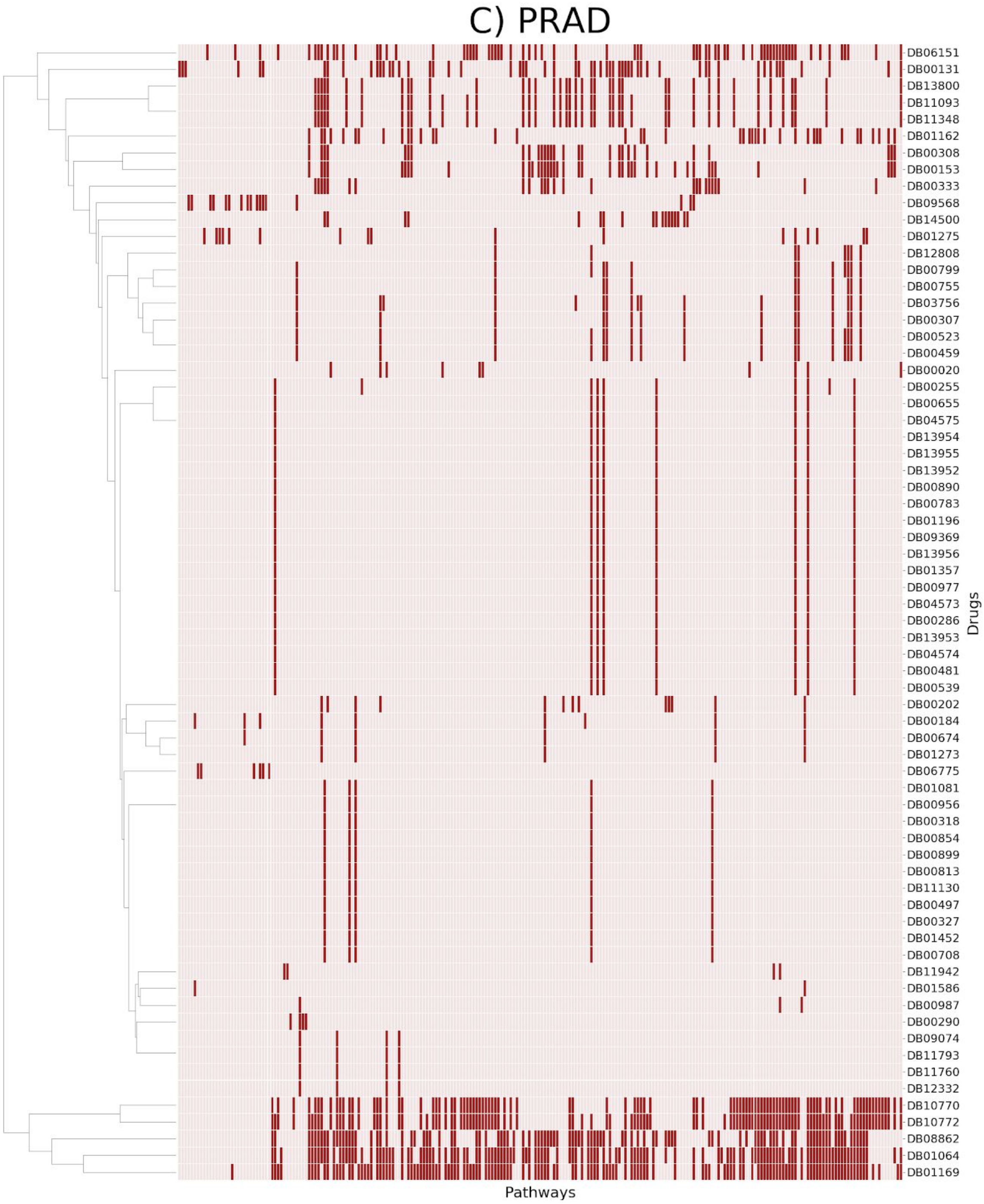


**Supplementary Figure 8. Pathways targeted by the prioritized drugs in DrugBank for PRAD.** The X-axis corresponds to the pathways targeted by any of the prioritized drugs (i.e., other KEGG pathways not targeted by any prioritized drug have been omitted for better visualization). Drugs (Y-axis) have been clustered based on the pathways they target. Due to the large number of pathways, we have clustered pathway groups together for visualization purposes. Black cells correspond to pathways targeted for each drug.
